# Efficient prestimulus network integration of FFA biases face perception during binocular rivalry

**DOI:** 10.1101/2021.06.25.449883

**Authors:** Elie Rassi, Andreas Wutz, Nicholas Peatfield, Nathan Wiesz

## Abstract

Ongoing fluctuations in neural excitability and connectivity influence whether or not a stimulus is seen. Do they also influence which stimulus is seen? We recorded magnetoencepahlography data while participants viewed face or house stimuli, either one at a time or under bi-stable conditions induced through binocular rivalry. Multivariate pattern analysis revealed common neural substrates for rivalrous vs. non-rivalrous stimuli with an additional delay of ~36ms for the bi-stable stimulus and poststimulus signals were source-localized to the fusiform face area (FFA). Prior to stimulus onset followed by a face-vs. house-report, FFA showed stronger connectivity to primary visual cortex and to the rest of the cortex in the alpha frequency range (8-13 Hz) but there were no differences in local oscillatory alpha power. The pre-stimulus connectivity metrics predicted the accuracy of post-stimulus decoding and the delay associated with rivalry disambiguation suggesting that perceptual content is shaped by ongoing neural network states.

## Introduction

Ongoing neural fluctuations interact with sensory inputs and influence perception (1, 2). These interactions can be captured by characterizing prestimulus brain activity and the influence of that activity on upcoming perception and behavior. For example, prestimulus oscillatory power, an index of neural excitability, influences whether or not a near-threshold stimulus will be detected (3–6). Considering the complex spatio-temporal properties of ongoing neural activity, connectivity measures have also been used to better understand prestimulus brain states. As a result, prestimulus connectivity was also shown to influence near-threshold perception (7–10).

However beyond mere stimulus detection, one of the most important functions of the visual system is categorizing objects thereby building the contents of visual experiences (11–13). To distill the neural correlates of visual content, researchers have employed bistable perception paradigms in which the perceived category of an ambiguous but invariant stimulus can switch over time (14). Examples of bi-stable stimuli include the Necker cube (15), the Rubin vase (16), and binocular rivalry (17). When bistable stimuli are presented briefly, only one of the two possible interpretations of the ambiguous image is perceived. Over multiple presentations, perception of the stimulus will vary between the two interpretations. This trial-by-trial variance in perception can then be used to assess the prestimulus patterns that might have influenced perception of one or the other stimulus category (18, 19). As such, briefly presented bi-stable images are unique tools to probe the influence of prestimulus neural activity on subsequent perception. Given that prestimulus neural excitability and connectivity bias stimulus detection (‘Did you see anything?’), here we ask whether they also bias perceptual content (‘ What did you see?’).

Previous studies that have used bi-stable stimuli with semantic content, such as the faces and the vase of the Rubin vase, have reported the involvement of temporal cortical regions known to be involved in non-rivalrous object perception. For example, an fMRI study using the Rubin vase illusion reported increased prestimulus Fusiform Face Area (FFA) activity prior to face (vs vase) reports (20), and a Magnetoencephalography (MEG) study using the same stimulus reported increased prestimulus connectivity between the FFA and early visual cortex prior to face (vs vase) reports (18). The connectivity patterns in this MEG study were found in the frequency range between 5 and 20 Hz covering theta, alpha, and lower beta oscillations. Among those, the effects were most pronounced in the alpha frequency range (~8-13 Hz). Alpha oscillations are the most prominent rhythm in the human brain and have been functionally linked with neural excitation and inhibition (21–23).

While the ambiguity of the Rubin vase illusion is likely resolved by means of figure-ground segregation (16, 24, 25), the perceptual processes involved in resolving binocular rivalry are less clear and still a matter of debate (26, 27). Even less known is the effect of ongoing prestimulus brain activity on resolving binocular rivalry, as the rivalrous stimulus is typically presented continuously to investigate the neural correlates of perceptual ‘switches’, or alternations between the two percepts (28, 29). To probe possible prestimulus effects at rivalry onset, the rivalrous stimulus would have to be presented briefly in between periods of no presentation. Such a set-up, in combination with the high temporal resolution of MEG, would further allow to explore the temporal dynamics of rivalry resolution or disambiguation, as compared to non-rivalrous perception.

Based on the “Windows to Consciousness” framework (10, 30), fluctuating connectivity levels influence upcoming perception and neural signals associated with perception. In particular, it states that the network integration of brain regions, which are crucial for mediating a particular perceptual experience (so-called “essential nodes”; (31)), will bias further processing, because more strongly integrated regions are more likely to broadly distribute its activity. The extent of network integration can be quantified using measures from graph theory such as efficiency (10, 32). In the present MEG study, we used a brief-presentation binocular rivalry set-up with the classical face and house combination. Treating the FFA as an essential node, we investigated whether prestimulus neural excitability and/or network integration of this region influences the disambiguation and perceptual report of the rivalrous stimuli, and whether the relevant prestimulus patterns also influence the corresponding post-stimulus neural activity.

## Results

### Behavioral reports are stochastic

We recorded whole-head MEG while 21 participants viewed an ambiguous image consisting of a red or blue face superimposed with a blue or red house. Participants wore red/blue glasses as we presented the image briefly and repeatedly, and asked them to report the content of their percept (face or house) on each trial. That is, we used the typical binocular rivalry setup but with short presentation times such that only one percept is formed, and asked participants to report that percept (Figure 1; bottom ‘Main Task’). Participants also passively viewed non-rivalrous versions of the face and house stimuli in a separate block prior to the main task (Figure 1; top ‘Passive viewing’).

**Fig. 1.**
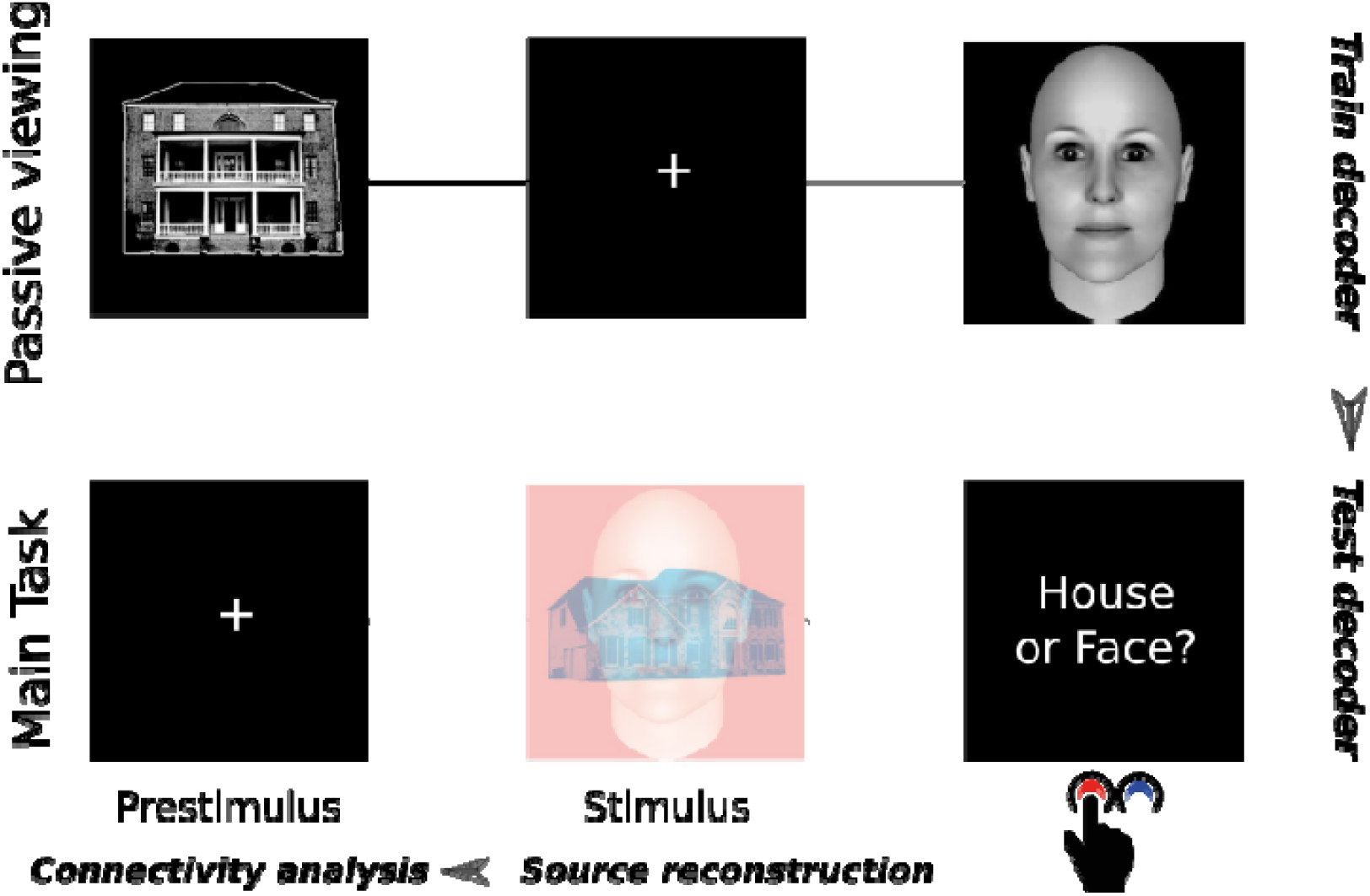
Experimental design. Example trial sequences from the passive viewing task where we trained the decoder (top) and the main experimental task where we tested the decoder (bottom).

To confirm that response choices in the main task were stochastic trial-by-trial, we binned consecutive responses in bins of 0 to 10 repetitions separately for face and house responses. We then averaged the number of repetitions in each bin across participants, and fit these averaged values with a binomial distribution. The distribution accounted well for both face and house repetitions (goodness-of-fit: *R*^2^ face = .92, *R*^2^ house = .89). In other words, participants were equally likely to report “face” or “house” on each trial, regardless of their responses on previous trials. Overall, face reports constituted 40% of reports. For all subsequent analyses, we equalized the face and house trial numbers per participant by randomly omitting trials from the pool with more trials.

### Rivalry resolution is delayed compared to non-rivalrous vision

To explore the temporal dynamics of rivalry resolution, we first investigated the differences and similarities in neural activity between perceiving a non-rivalrous face or house image and perceiving a face or house under binocular rivalry. For this purpose, we also recorded MEG while participants passively viewed a series of unambiguous, non-rivalrous face and house images prior to the main experiment. The motivation was twofold. First, we wanted to be confident that participants actually reported what they had perceived. Second, we wanted to investigate the time course for non-rivalrous vs. rivalrous images. To achieve this, we used linear discriminant analysis to train a classifier on the broadband MEG signals resulting from the non-rivalrous images, and tested it on the signals resulting from the rivalrous images using the temporal generalization method (33).

The classifier performed significantly above chance in the post-stimulus period (0 to 500 ms; p(cluster)<.001), meaning that the non-rivalrous and rivalrous conditions shared decodable neural representations (Fig 2a). The signals from the non-rivalrous data at 85ms onwards could significantly predict those of the rivalrous data from 120 ms onwards. Decoding accuracy peaked outside of the diagonal of the time generalization matrix, indicating a delay in decodability. To quantify this delay, we calculated the difference between the time at which the non-rivalrous data could most predict the rivalrous data, and the time at which the rivalrous data could most predict the non-rivalrous data (Fig 2b). The difference between these two latencies was on average 36 ms and it was significant (T=6.8, p<.0001). Overall, this analysis gave us confidence that participants truly reported what they had perceived, and showed that visual content is decodable from brain responses to ambiguous and unambiguous stimuli. More importantly, this result indicates that when faced with a bistable face-house stimulus the brain requires approximately an additional 36 ms to resolve this ambiguity.

**Fig. 2.**
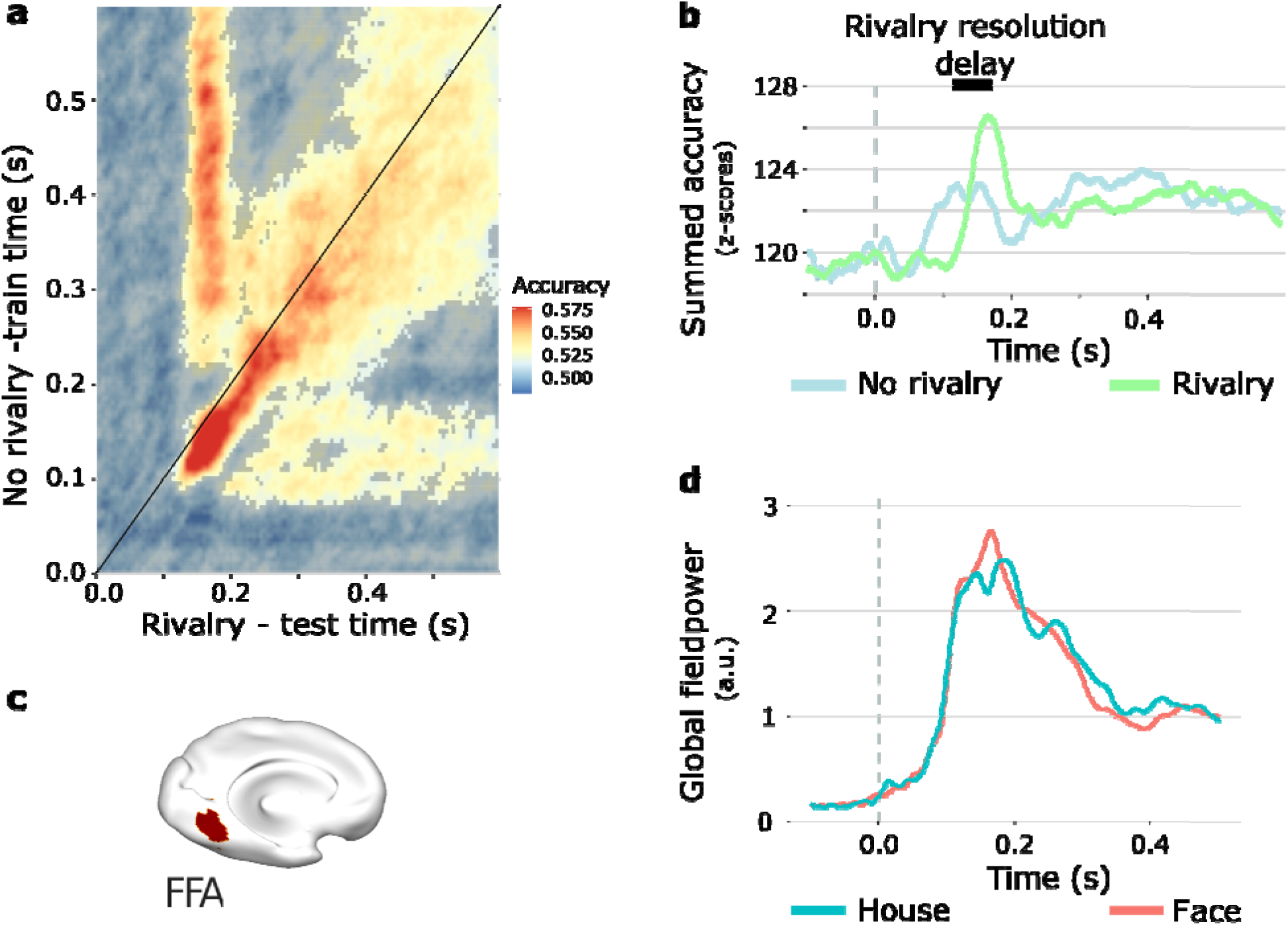
Post-stimulus responses to rivalrous and non-rivalrous face/house images. **(A)** Face vs house decoding of sensor-level data using the time-by-time generalisation method, obtained from training a classifier on evoked responses to unambiguous face and house images, and testing on evoked responses to the bistable face/house images. (B) Peak decoding accuracy of the rivalrous data was around 36 ms later than the time of peak decoding accuracy of the non-rivalrous data. (C) Source-localised region of interest: FFA. (D) Global field power in source space (averaged across all voxels).

To answer our main question of whether prestimulus activity in category-sensitive brain regions biases upcoming perceptual content, we first defined such a region in a data-driven manner. We used a beamforming approach to project our data to source space and grand-averaged the evoked responses to faces and houses across sources and participants (Fig 2d). The difference between the two evoked responses was significant during the post-stimulus period (0 to 500 ms; p(cluster)<.01). This difference peaked 165 ms after stimulus onset, in a similar time window in which the MVPA classifier achieved peak decodability. We then defined our region of interest as the region where this difference was strongest. Consistent with previous literature (34), we localized the difference in responses to faces and houses in the left fusiform gyrus (Fig 2c; FFA; MNI coordinates: [−28, −58, −8] mm). For the subsequent analyses, we used FFA as our region of interest.

### Prestimulus connectivity biases perceptual report

Next, we investigated whether oscillatory power and connectivity in FFA was different before the participants reported “face” vs “house”. We used a non-parametric cluster-based permutation approach (35) to correct for multiple comparisons over the 5 to 20 Hz frequency range. The frequency spectra from both trial types showed that oscillatory activity was largely restricted to the 5 to 20 Hz range, with clear alpha band peaks around 10 Hz. However, we found no statistical differences in prestimulus power between face and house trials (Fig 3a). This finding is consistent with a recent similar study similar, which reported increased connectivity between FFA and primary visual cortex (V1) prior to face reports, despite no differences in power (18). We therefore additionally investigated the prestimulus coherence between FFA and atlas-defined V1. We calculated coherence in a frequency-resolved manner and used a cluster-based permutation approach to contrast prestimulus coherence between FFA and V1 on face vs house trials. Here we restricted our statistical testing to the frequency range in which the aforementioned study reported the coherence effect (10-16 Hz). We found increased prestimulus coherence in the alpha range on face (vs house) trials (p(cluster)=0.021; Fig 3b). Consistent with the aforementioned study, FFA and V1 were more strongly connected prior to face (vs house) responses, despite no differences in power between the two trial types. Crucially, this meant that connectivity effects would not be confounded with power differences in the current dataset.

**Fig. 3.**
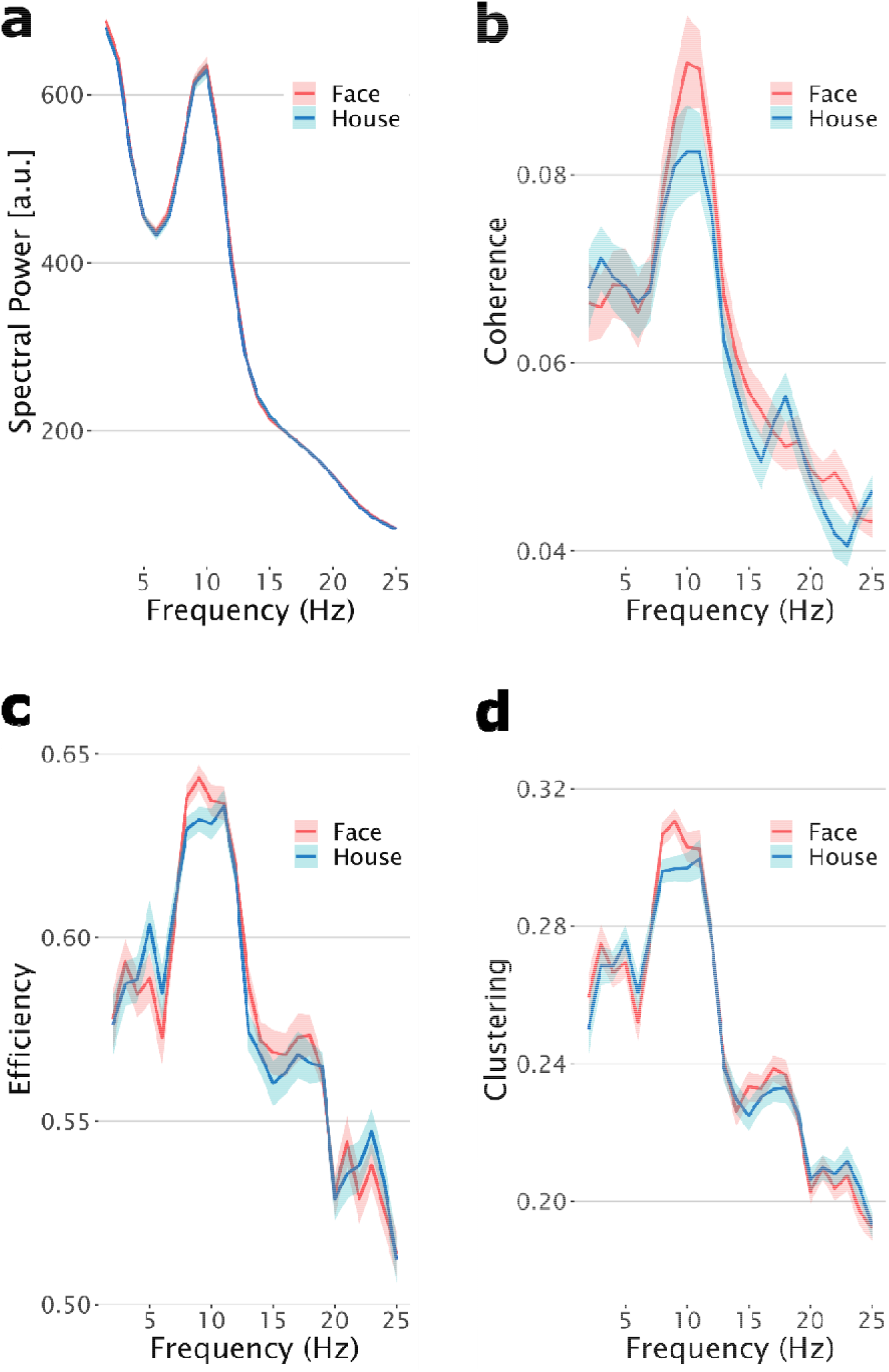
Prestimulus MEG connectivity biases upcoming perception. Shaded areas represent the standard error of the mean. (A) No difference in prestimulus spectral power between face and house trials. (B) Prestimulus coherence between V1 and FFA was increased on face trials compared to house trials. **(C)** Prestimulus efficiency was increased on face trials compared to house trials. **(D)** Prestimulus clustering was increased on face trials compared to house trials.

**Fig. 4.**
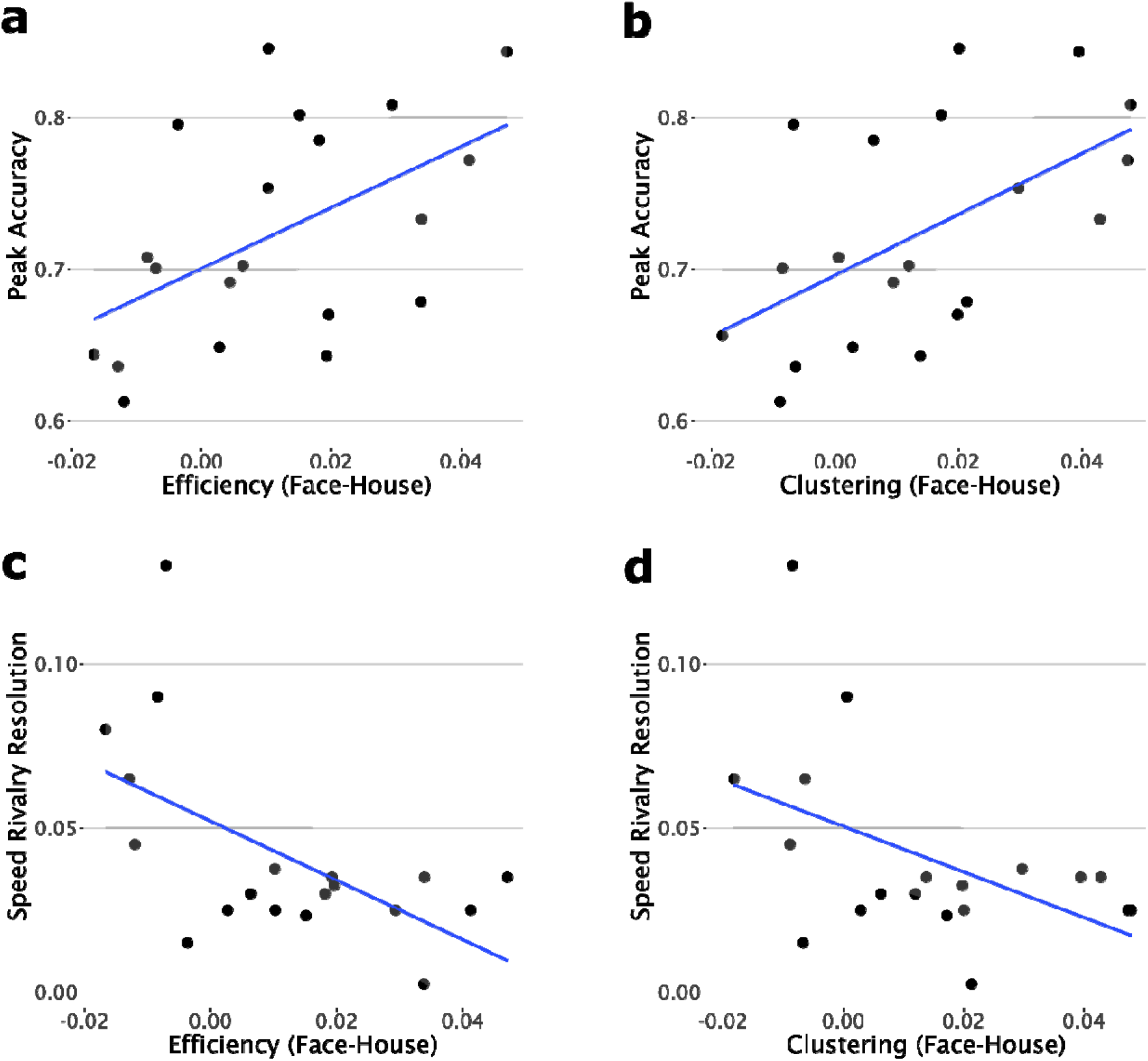
Prestimulus connectivity is correlated with post-stimulus decoding accuracy and rivalry resolution across participants. r values represent Pearson’s correlation coefficients. Shaded areas represent 95% confidence intervals. (**A**) Mean prestimulus efficiency differences are correlated with maximum post-stimulus decoding accuracy. (**B**) Mean prestimulus clustering differences are correlated with maximum post-stimulus decoding accuracy. (**C**) Mean prestimulus efficiency differences are negatively correlated with speed of rivalry resolution. (**D**) Mean prestimulus clustering differences are negatively correlated with speed of rivalry resolution.

In the next step, we investigated whether the network integration patterns of FFA influence the perceptual reports. To assess this information, we applied two local graph theoretical measures to thresholded (i.e. binary) connectivity matrices. The first was ‘local efficiency’, a measure of how well-integrated a node is to the rest of the network estimated by the inverse of the average shortest path from this node to all other network nodes. The second was the ‘local clustering coefficient’, a measure of the robustness of the region’s local network (i.e. the probability that nodes to which the region is connected are connected among each other). We computed these measures in a frequency-resolved manner, and used a cluster-based permutation approach (clustering over the frequencies 5-20 Hz) to contrast prestimulus efficiency and clustering of FFA between face and house trials. We found that FFA showed increased prestimulus efficiency and clustering on face trials compared to house trials (Fig 3c, efficiency: p(cluster) = 0.017; Fig 3d, clustering: p(cluster) = 0.024), despite no differences in spectral power. In both cases, the difference was most pronounced in the alpha band, particularly at frequencies 8-9 Hz. In other words, FFA was more efficiently connected to the rest of the brain in a robust manner prior to face (vs house) responses.

### Prestimulus connectivity predicts post-stimulus decoding accuracy and rivalry resolution

Finally, we investigated whether prestimulus connectivity predicts post-stimulus category-related and rivalry-related neural activity across participants. For each participant, we extracted the maximum face vs house decoding accuracy and the average rivalry-related delay in decodability from the time generalisation results, as well as the average differences between face and house trials in prestimulus efficiency and clustering. Maximum decoding accuracy was positively correlated with both the prestimulus efficiency (r=.55, p=.010) and prestimulus clustering (r=.59, p=.005) differences. In other words, participants who showed stronger prestimulus connectivity effects also showed more robust post-stimulus decoding. Consequently in addition to being predictive of upcoming perceptual reports, prestimulus connectivity was also predictive of the strength of post-stimulus signals associated with those percepts.

Further, the rivalry-related delay in post-stimulus decoding was negatively correlated with both the prestimulus efficiency (r=-.50, p=.022) and prestimulus clustering (r=-.53, p=.013) differences. In other words, participants who showed stronger prestimulus effects also had shorter delays in post-stimulus rivalry decodability. Overall, these results indicate that the influence of prestimulus connectivity on post-stimulus neural signals is manifested in higher decoding accuracy and shorter delays in rivalry resolution.

## Discussion

While the prestimulus requisites of near-threshold conscious perception have been widely studied (1–10), very few studies have investigated the influence of prestimulus activity on object perception (20, 36). These fMRI studies reported increased activity in category-selective extrastriate regions prior to object perception, but these prestimulus BOLD activations could represent increased local excitability of a cortical region, or they could be a product of increased connectedness of that region. In fact, a recent MEG study has reported that increased prestimulus connectivity between striate and extrastriate regions, but not local excitability of these regions, predicted upcoming object perception (18), a finding that we replicate in the current study.

Here we aimed to add to this emerging literature by employing a paradigm similar to the one in the aforementioned MEG study and investigating possible influences of prestimulus excitability and connectivity on object perception in a binocular rivalry setup. We showed people a bi-stable face/house image whose perceptual content would vary trial-by-trial, and asked them to report their percept on each trial. We source-localized the post-stimulus activity to FFA and subsequently focused on differences between face-report and house-report trials in prestimulus FFA activity. We found no differences in FFA oscillatory alpha-band power prior to trials perceived as face vs house. Nevertheless, we did observe differences in FFA connectivity states prior to trials perceived as face vs house. Specifically, FFA was more efficiently and robustly connected to the rest of cortex in the alpha frequency range prior to face (vs house) perception of the bi-stable face/house image. These connectivity differences were correlated with post-stimulus decoding accuracy and speed of rivalry resolution across subjects.

Alpha oscillations are implicated in a variety of cognitive and perceptual phenomena (37) and act on neuronal populations by rhythmic inhibition (38). Their power can therefore be thought of as a proxy to neural excitability. If the prestimulus levels of excitation per se of the task-relevant sensory cortices influenced upcoming object perception, we would expect prestimulus alpha power in FFA to be different on face vs house trials. Yet we did not observe such a difference. Despite this, we found differences between face and house trials in the prestimulus connectivity states of FFA. While prestimulus excitability of task-relevant sensory cortices influences simple perceptual operations such as near-threshold detection (39, 40), our findings indicate that the connectivity states of these task-relevant regions are more relevant to understanding more complex perceptual operations, such as visual object perception.

Graph theoretical measures of network connectivity have provided important insights into properties of structural and functional brain networks, such as small-worldness and the existence of hubs (32, 41). Here we used the graph theoretical measures of efficiency and clustering coefficient to inform us about the connectivity states of FFA prior to face and house perception. We found increased prestimulus local efficiency in FFA on face (vs house) trials. This means that FFA was more highly integrated into the rest of the network prior to face (vs house) perception. In other words, when FFA was better integrated in cortex, people were more likely to subsequently perceive a face rather than a house. We additionally found increased prestimulus local clustering in FFA on face (vs house) trials. This means that FFA was more densely connected to its neighbors prior to face (vs house) perception. In other words, when the network to which FFA belongs was more robust, participants were more likely to subsequently perceive a face rather than a house.

Our results provide further support to the Windows to Consciousness framework (10, 30), which postulates that ongoing neural fluctuations impact pre-established connectivity pathways and influence upcoming perception. We now widen this view to include object perception, which is known to involve ventral stream activity, especially in the case of stimuli with semantic content. Since our time generalization decoding analysis supported the notion that ambiguous and unambiguous perception share common neural substrates, we hypothesized that the ventral visual pathway be the candidate pre-established connectivity pathway. In support of this hypothesis, we found increased coherence between V1 and FFA prior to face (vs house) perception.

Despite the similarity in neural responses to rivalrous and non-rivalrous images, we found an important difference between them. Decoding of responses to the rivalrous images was delayed with respect to those of the non-rivalrous images. This possibly relates to a delay in disambiguating the bistable image, before perceptual content becomes consciously accessible. In other words, this delay can be thought of as representing the speed at which rivalry is resolved on average within a participant. In fact, this delay was negatively correlated with the prestimulus connectivity effects, indicating that participants showing stronger connectivity effects also resolved the binocular rivalry faster. Finally, we reasoned that if prestimulus connectivity biases upcoming object perception, then it should also bias the post-stimulus signals associated with object perception. We found that both prestimulus efficiency and clustering correlated well with face vs house decoding accuracy scores. The stronger the prestimulus effect was, the more decodable the post-stimulus signals were. This could indicate that participants who were more prone to the prestimulus connectivity effects were also better at subsequently disambiguating the bistable stimulus. Overall, it appears that prestimulus connectivity biases not only perceptual contents, but also the strength and speed of content-related neural activity.

It is difficult to assign cognitive functions to the connectivity effects we detected, but at least two interpretations of these effects are plausible. One is that a conscious, topdown drive to perceive one of the contents of the stimuli leads to increased connectivity of the relevant cortical region, which in turn biases perception. This interpretation is in line with a large body of literature suggesting that expectations, predictions, or context critically shape perception (42–45). That the effects were in the alpha frequencies might also support this interpretation, given the evidence from human and non-human primate studies showing that alpha oscillations subserve feedback connectivity in visual cortical areas (46, 47). However, a purely top-down interpretation is unlikely given our design and behavioral analysis: the prestimulus intervals were short (1 to 2 sec) and stimulus onset was jittered and difficult to predict within this range, and in addition, our behavioral analysis indicated that the response sequences were stochastic. Therefore, the other plausible interpretation is that spontaneous fluctuations in neural activity, which could randomly differ on a trial-by-trial basis, influence the connectivity state of the relevant cortical region, and in turn bias perception.

## Materials and Methods

### Experimental design

The main aim of the study was to test whether and how prestimulus brain activity recorded with MEG influences object perception. More specifically, we aimed to test whether perceiving a bi-stable face/house visual stimulus as “face” or “house” was biased by the prestimulus network states of category-selective brain regions. Additionally, we aimed to test how the neural activity associated with perceiving the bistable stimulus related to that associated with perceiving unambiguous stimuli. For these purposes, we employed a brief-presentation onset rivalry design, as well as a passive viewing task (oddball design) and staircase procedure. We describe the task designs in more detail below.

### Participants

21 volunteers with normal or corrected-to-normal vision participated in this experiment (11 m/10 f, 4 left-handed, mean age 26.3). During the course of the experiment participants wore non-magnetic clothes, and a questionnaire prior to the experiment excluded any metal artefacts on the participants being. The Ethics Committee of the University of Trento approved the experimental procedure and all participants gave written informed consent before taking part in the study.

### Experimental Procedure

Participants were seated upright in the MEG system and we instructed them to keep fixation, and to avoid eye blinks and movements as best as possible during the experiment. To create rivalry, we used Blue/Red Filter Glasses which participants wore throughout. We took into account the color filters and as such adapted the stimulus to suit the glasses. In between the blocks participants had a short break but remained seated in the MEG system. We displayed visual stimuli via a video projector outside of the MEG chamber which projected to a screen in the MEG chamber. A camera allowed us to monitor participants during the experiment.

The experiment was divided into three different task types: staircase thresholding, passive viewing/oddball task, and onset binocular rivalry. We programmed and controlled all experiments in Psychtoolbox (48) and the DataPixx I/O Controller and backprojected using the ProPixx Projector at 120Hz at 1980×1080 with the central images 400 x 400 in size and presented at a distance of 78cm.

### Staircase thresholding

Due to the competing face/house stimulus properties, we wanted to ensure that participants perceived roughly equal amounts of houses and faces. As such we ran a staircase thresholding procedure with the Matlab toolbox Palamedes (49). Four staircase procedures controlled the brightness of the red house, blue house, red face, and blue face stimuli. The procedure was a simple 1 up 1 down with a step-size of 0.01, a start value of 0.5, a min/max of 0.95/0.05, and a stop-rule at 32.

### Passive viewing/oddball task

To compare the neural responses to rivalrous and unambiguous stimuli, we presented participants with unambiguous house and face images (80 of each) prior to the main experiment. A stimulus would appear for 500 ms, followed by a jittered inter-stimulus interval of 1 to 2 sec, during which a fixation cross would appear. We required participants to report an oddball (inverted house or face) occuring on 10% of the trials to keep them engaged in the viewing task.

### Onset binocular rivalry (main task)

In the main experiment we presented participants with the rivalrous face/house images and required them to report their percept on each trial. A trial would begin with the presentation of a central fixation cross, followed by a blank screen for jittered interval of 1 to 2 sec. After the blank screen the bistable image would appear for 500 ms, and then a response screen ‘House or Face?’ would appear, the prompt for participants to respond with a right-handed button press to indicate their percept. The experiment consisted of 6 blocks of 80 trials each (total of 480 trials).

### MEG Data Acquisition

We recorded MEG using a 306 channel whole head VectorView MEG system (Elekta-Neuromag, Ltd., Helsinki, Finland, 204 gradiometers and 102 magnetometers) installed in a magnetically shielded chamber (AK3b, Vakuumschmelze Hanau, Germany), with signals recorded at 1000 Hz sampling rate. We adjusted the hardware filters to bandpass the MEG signal in the frequency range of 0.01 Hz to 330 Hz. Prior to the recording, we recorded points on the participant’s head using a digitizer (Polhemus, VT, USA). These points included the 5 HPI coils, the three fiducials (nasion, left and right pre-auricular points), and around 300 additional points covering the head as evenly as possible. We used the HPI coils to monitor head position during the experiment.

### MEG Preprocessing and Source Projection

We pre-processed the data using the Fieldtrip toolbox (50). From the raw continuous data, we extracted epochs of 4 seconds lasting from 2.5 seconds before onset of the picture to 1.5 seconds after stimulus onset. This resulted in 400 trials per participant. We applied a high-pass filter on this epoched data at 1 Hz (IIR Butterworth 6-order two-pass filter with 36 dB/oct roll-off), followed by a band-stop filter of 49-51Hz to remove power line noise. We then down-sampled the data to 400 Hz. We visually inspected the trials for strong physiological (e.g. blinks) and non-physiological artefacts (e.g. channel jumps) and rejected the contaminated trials. For each participant, we then assigned the trials to the 2 conditions according to the participants’ responses. Finally we equalized the number of trials in each condition to ensure comparable signal-to-noise ratio, which left us with 176 trials per condition on average.

We projected the data to source space by applying LCMV (linear constrained mean variance) beamformer filters to the sensor level data (51). To create anatomically realistic headmodels (52), we used structural MRI scans and the Polhemus digitized scalp shape. We used individual MRI scans for the 5 participants for which the scans were available, and for the rest, we used a template MRI which was morphed to fit the individuals head shape using an affine transformation. We calculated a threedimensional source grid (resolution: 8 mm) covering an entire MNI standard brain volume. For each of these points, we computed an LCMV filter using the individual leadfield and the data covariance matrix. We then multiplied the sensor-level signal to the LCMV filters to obtain the time series for each source location in the grid and for each trial. Finally, we used a parcellation scheme of 333 parcels (53) and averaged activity within each parcel.

To localize our region of interest, we first calculated the time-locked averages of the parcellated source-level signals (band-pass filtered between 1 and 30 Hz), normalized the responses across participants, and baseline-corrected them with a relative baseline of −200 ms to 0. We then computed the difference in the grand-averaged evoked responses between face and house trials (across participants and sources) to determine the time point at which there was the largest difference, which was at 165 ms. Finally, we contrasted face and house evoked response amplitude at that time point across participants with t-statistics for each parcel source and chose the parcel with the largest t-stat as our region of interest.

### Multivariate Pattern Analysis (MVPA) decoding

We performed MVPA decoding on the post-stimulus period 0-500 ms based on normalized (z-scored) single-trial sensor data downsampled to 100Hz and band-pass filtered between 0.1 Hz and 40 Hz. We used multivariate pattern analysis as implemented in CoSMoMVPA (54) to identify when a common network between perception of unambiguous and rivalrous stimuli was activated. The two classes were face and house; in the rivalrous case,we labelled trials as face and house according to the participants’ responses. The training set were the unambiguous trials and the testing set were the rivalrous trials. We used a linear discriminant analysis (LDA) classifier with the temporal generalization method (33) to assess the ability of the classifier across different time points in the training set to generalise to every time point in the testing set. We used local neighborhoods features in time (in each time step of 10 ms we included as additional features the previous and following time steps). We generated the time generalization matrix for each participant and statistically tested for above-chance decoding (defined as >50% accuracy) significance with a cluster-based permutation approach (clustering over the two time dimensions between 0-500 ms).

To calculate the difference in peak decoding latencies between training and testing data, we averaged the time-by-time spectrum across each of its two dimensions and identified the two timepoints at which these traces peaked, before subtracting them.

### Analysis of prestimulus power, graph measures, and coherence

We calculated spectral power, efficiency, the clustering coefficient in source-localized left FFA, as well as coherence between FFA and left V1 (defined by the parcellation scheme), in the prestimulus interval of −1 to 0 sec. We used multi-taper frequency transformation with a spectral smoothing of 2 Hz to get Fourier coefficients. From those we extracted power in FFA and computed coherence between FFA and V1.

For the graph analysis we computed the cross-spectral density (CSD) on source data and derived an all-to-all connectivity matrix of imaginary coherence, a conservative measure of phase synchronization. We then took the absolute values of the connectivity matrix calculated the adjacency matrix by thresholding the connectivity matrix. We set the thresholds by taking the maximum value across the first dimension of the (source x source x frequency x time) connectivity matrix, and then the minimum values of those maximum values across the 2nd dimension of the matrix, obtaining a threshold per frequency and time-point. Finally, we used the connectivity matrix and thresholded adjacency matrix to calculate local efficiency and local clustering coefficient in FFA. We averaged all of the above measures over the prestimulus time interval, and contrasted face vs house estimates across participants with a cluster-based permutation approach.

### Statistical Analysis

For all statistical analyses, we used nonparametric cluster permutation tests (35), comparing a selected test statistic against a distribution obtained from 10000 permutations. We set thresholds for forming clusters at p <.05 and considered an effect significant if its probability with respect to the nonparametric distribution was p < .05. For the evoked responses, we used 2-sided dependent-samples T-tests and clustered across the time dimension (0 to 500 ms). For the MVPA data, we used 1-sided T-tests (as we were interested in above-chance decoding) and clustered across the two time dimensions (both 0 to 500 ms). For the spectral power, efficiency, and clustering measures, we used 1-sided T-tests (as we hypothesized greater values for “face” trials) and clustered across the frequencies 5 to 20 Hz. For the coherence measure, we did the same but clustered over the frequencies 10 to 16 Hz. For the correlations, we computed Pearson’s coefficients.

## Acknowledgments

We would like to thank Anne Hauswald, Chrysa Lithari, and Gaëtan Sanchez for assistance and valuable input on data analysis.

## Author contributions

Conceptualization: ER, NP, NW

Methodology: ER, NP, NW

Investigation: ER, NP

Visualization: ER

Supervision: AW, NP, NW

Writing—original draft: ER

Writing—review & editing: AW, ER, NP, NW

## Data and materials availability

The data and code used in the analysis will be made publicly available on the Open Science Framework (OSF) server upon acceptance of the manuscript for publication.

